# Simulation of Energy Regeneration in Human Locomotion for Efficient Exoskeleton Actuation

**DOI:** 10.1101/2022.06.13.495983

**Authors:** Brokoslaw Laschowski, Keaton A. Inkol, Alex Mihailidis, John McPhee

## Abstract

Backdriveable actuators with energy regeneration can improve the efficiency and extend the battery-powered operating times of robotic lower-limb exoskeletons by converting some of the otherwise dissipated energy during negative mechanical work into electrical energy. However, previous related studies have focused on steady-state level-ground walking. To better encompass real-world community mobility, here we developed a feedforward human-exoskeleton energy regeneration system model to simulate energy regeneration and storage during other daily locomotor activities. Data from inverse dynamics analyses of 10 healthy young adults walking at variable speeds and slopes were used to calculate the negative joint mechanical power and work (i.e., the mechanical energy theoretically available for electrical energy regeneration). These human joint mechanical energetics were then used to simulate backdriving a robotic exoskeleton and regenerating energy. An empirical characterization of the exoskeleton device was carried out using a joint dynamometer system and an electromechanical motor model to calculate the actuator efficiency and to simulate energy regeneration. Our performance calculations showed that regenerating energy at slower walking speeds and decline slopes could significantly extend the battery-powered operating times of robotic lower-limb exoskeletons (i.e., up to 99% increase in total number of steps), therein improving locomotor efficiency.

## I. Introduction

Robotic lower-limb exoskeletons can replace the propulsive function of impaired biological muscles and allow persons with mobility impairments, due to aging and/or physical disabilities, to perform daily locomotor activities that require power generation [1]. Most exoskeletons use electromagnetic actuators for power generation, specifically brushed or brushless direct current (DC) motors [1]. These electric motors are often coupled with a high-ratio transmission (e.g., ball-screw mechanism or harmonic gearing) to increase the motor torque output to that needed for human locomotion, which causes the robotic actuator to have high output impedance [2]. For example, the ReWalk and Ekso Bionics powered lower-limb exoskeletons use stiff actuators to rigidly track predefined kinematic trajectories for precise position control, which can benefit users with limited ability to physically interact with and control the robotic device (e.g., persons with complete spinal cord injury) [1].

However, these traditional stiff actuators cannot exploit the passive dynamics of human locomotion and/or other energy storage and return mechanisms, therein often resulting in heavy and inefficient actuators that require significant energy consumption and thus provide limited battery-powered operating times given the finite energy density of rechargeable batteries. Most robotic lower-limb exoskeletons provide only 1-5 hours of maximum battery-powered runtime [1]. Onboard portable power has often been considered one of the leading challenges to developing robotic exoskeletons for real-world mobility [1]. Increased device mass and inertia, due to heavier onboard motors and batteries, would also require more effort by the human musculoskeletal system during swing phase, thus reducing the locomotor efficiency via higher metabolic power consumption [3]. To address these limitations, researchers have been working on designing lightweight and efficient actuators that more effectively utilize the energetics of human locomotion.

Torque-dense motors with low-ratio transmissions, known as quasi-direct drives, have been used to achieve low mechanical impedance and high backdrivability and efficiency [4]. The use of low transmission ratios has largely been driven by recent advances in torque-dense motors in the drone industry. These actuators generate high output torque by increasing the motor torque density rather than the transmission ratio, thus reducing the effects of high gearing – i.e., increased weight, backlash, and reflected inertia, which scales with the transmission ratio squared [2]. Gears also have torque-dependent friction, which further increase the impedance and reduce backdrivability and efficiency [5]. High external loads are thus needed to overcome the impedance to backdrive the actuator. These characteristics of high gearing can impede dynamic physical interactions between the human and device and between the device and environment, which could especially encumber those with partial motor control function (e.g., older adults and/or persons with osteoarthritis or poststroke) who may benefit from the ability to actively backdrive the joints.

Backdriveable actuators with low output impedance have many benefits for control and efficiency, including: 1) free-swinging dynamic leg motion, which can simplify the control during swing phase and allow for more energy-efficient locomotion; 2) compliant impacts; 3) negligible unmodeled actuator dynamics, which can further simplify the control; 4) intrinsic backdriveability; and 5) energy regeneration during negative mechanical work [4]. Energy regeneration is the process of converting some of the otherwise dissipated energy during negative mechanical work into electrical energy via backdriving the actuator. In other words, when backdriven by an external load, the motor can provide a braking torque to decelerate the load (e.g., motion control during swing phase) while concurrently generating electricity [6]. This is analogous to regenerative braking in electric and hybrid electric vehicles. Assuming an adequate motor driver, the regenerated energy could be used for battery recharging and/or transferred to other joints to support power generation. These design principles were used in the MIT Cheetah robot [4] and more recently in the wearable robotic devices by Gregg and colleagues [7]–[10].

Although regenerative actuators can improve exoskeleton efficiency and extend the battery-powered operating times or decrease the weight of the onboard batteries, previous studies have focused on steady-state level-ground walking [9], [11], [12]. However, real-world community mobility involves variable activities like walking at different speeds and inclines as well as transitions between locomotion modes [13]. Non-steady conditions such as transitions between locomotion modes and variations in slopes and speeds are critical to modeling human locomotion and designing wearable robotic devices for real-world mobility. This observation is supported by our recent development of the *ExoNet* dataset, which showed that a relatively small percent (∼8%) of real-world walking environments consist of only continuous level-ground terrain [14], [15]. Motivated to explore energy regeneration during other daily locomotor activities, here we developed a feedforward human-exoskeleton energy regeneration system model to simulate backdriving a robotic exoskeleton with human joint mechanical power data while walking at variable speeds and slopes. We performed benchtop testing with the exoskeleton device using a joint dynamometer system [16], which, together with an electromechanical motor model, was used to calculate the actuator efficiency and to simulate energy regeneration and storage during human locomotion.

## II. Methods

### A. Energetics of Human Locomotion

We used the open-source biomechanics dataset recently published by [17], which was created to aid the development of biomechanical models of human locomotion and the design and control of wearable robotic devices; their study was approved by the review boards at the University of Texas at Dallas and the University of Michigan. The dataset includes hip, knee, and ankle joint mechanical powers of ten able-bodied subjects (age: 30 ± 15 years; height: 1.73 ± 0.94 m; weight: 74.6 ± 9.7 kg) walking at variable speeds (0.8 m/s, 1 m/s, and 1.2 m/s) and slopes (0° and ± 5° and 10°). 3D kinematics and ground reaction forces were measured using an optical motion capture system (Vicon, 100 Hz) and a Bertec instrumented split-belt treadmill, respectively. Joint powers in the sagittal plane were calculated from rigid-body inverse dynamics (i.e., the dot product of the net joint torque and angular velocity) 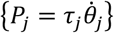. The joint mechanical power outputs (W/kg) were normalized to total body mass and percent stride (0-100%) to allow for between and within subject averaging, and interpolated (i.e., heel-strike to heel-strike) to have the same length (Fig. 1 and 2).

**Figure 1.**
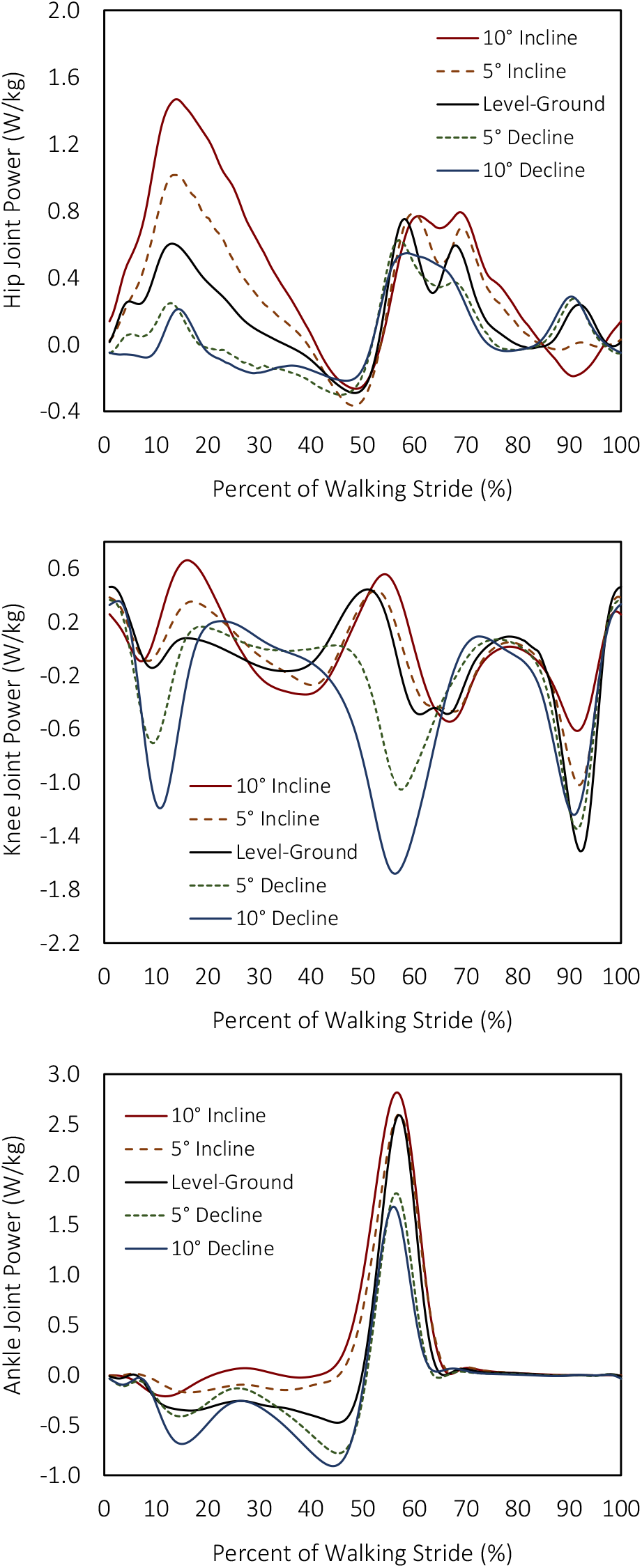
Hip, knee, and ankle joint mechanical power (W/kg) per stride in healthy young adults walking at 1 m/s on variable slopes (0° and ± 5° and 10°). Results are normalized to total body mass and percent of stride. The positive and negative values represent joint power generation and absorption, respectively. Data were calculated from [17], the trajectories of which begin and end with heel-strike. The results are averages across multiple subjects (n=10) and strides.

**Figure 2.**
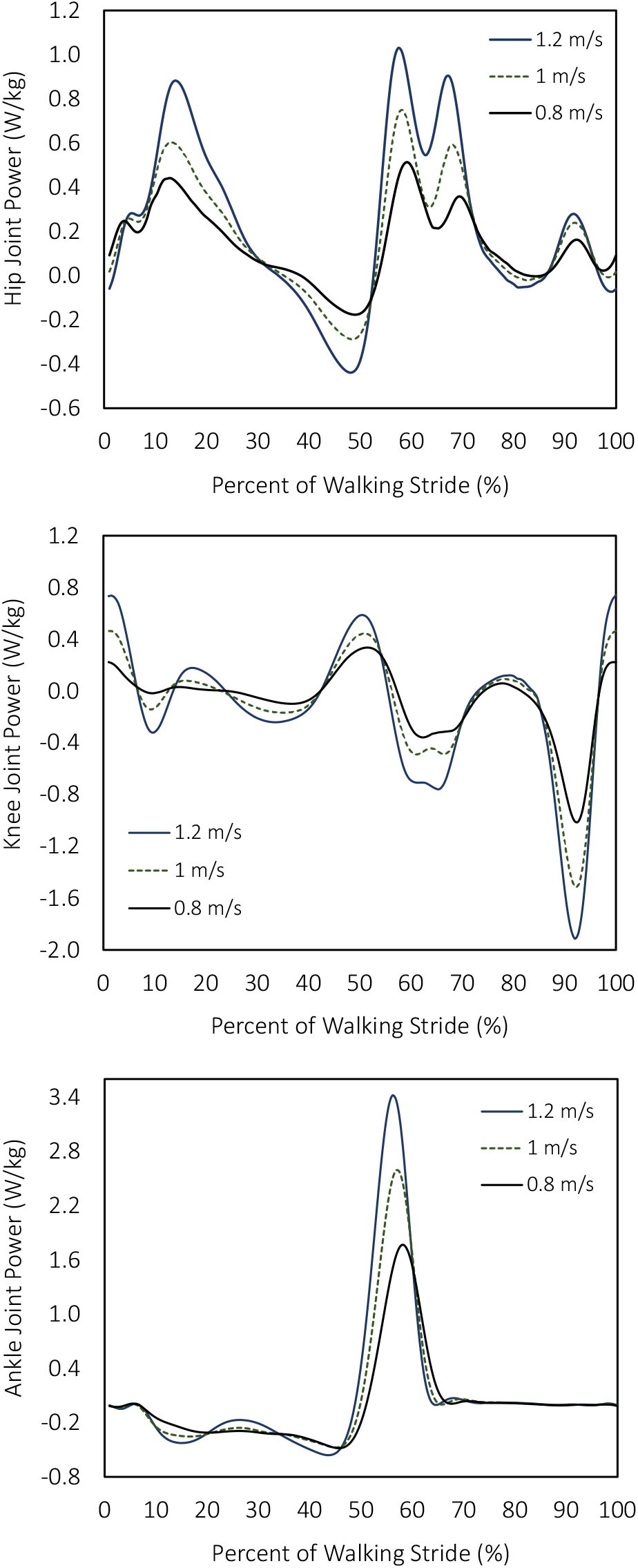
Hip, knee, and ankle joint mechanical power (W/kg) per stride in healthy young adults walking on level ground at variable speeds (0.8 m/s, 1 m/s, and 1.2 m/s). Results are normalized to total body mass and percent of stride. The positive and negative values represent joint power generation and absorption, respectively. Data were calculated from [17], the trajectories of which begin and end with heel-strike. The results are averages across multiple subjects (n=10) and strides.

We integrated the joint mechanical powers over time to calculate the joint mechanical energy generated and absorbed during human locomotion (Appendix 1). During energy generation, the net joint torque and angular velocity have the same sign direction and positive mechanical work is done (e.g., a concentric contraction wherein the muscles shorten under tension). During energy absorption, the net joint torque and angular velocity have opposite polarities and negative mechanical work is done (e.g., an eccentric contraction wherein the muscles lengthen under tension); this assumes that the joint torque generators are independent of adjacent joints such that biarticulating muscles spanning multiple joints are ignored. The net rate of energy generation or absorption by all muscles crossing the joint is the joint mechanical power. Note that muscle work, not necessarily joint work, is associated with the metabolic energetics of human movement. Therefore, the design and control of a regenerative actuation system based on joint mechanical work and power alone could bring about a metabolic penalty such that the net joint work is negative but some muscles crossing the joint could be doing positive work.

Figs. 1 and 2 show the calculated hip, knee, and ankle joint mechanical powers during human locomotion at variable speeds and slopes, respectively. Generally speaking, the knee behaves like a damper mechanism, performing net negative mechanical work with four main power phases: 1) negative mechanical power absorption at weight acceptance wherein the knee flexes under the control of an extensor moment; 2) positive mechanical power generation by the knee extensors during mid-stance such that the product of the extensor moment and angular velocity is positive; 3) negative mechanical power absorption by the extensors as the knee flexes during early swing; and 4) negative mechanical power absorption by the knee flexors during late swing to decelerate leg extension prior to heel-strike. In contrast, the ankle generally behaves like an actuating motor, performing net positive mechanical work with two main power phases: 1) negative mechanical power absorption at weight acceptance wherein the product of the plantarflexor moment and dorsiflexor velocity is negative; and 2) a significant positive mechanical power burst by the plantarflexors during push-off. The hip joint power is relatively small and irregular.

The phases of negative joint mechanical work during human locomotion present an opportunity to improve exoskeleton efficiency by recycling some of the otherwise dissipated energy using backdriveable actuators with energy regeneration. Next, we used these human joint mechanical energetics to simulate backdriving an exoskeleton efficiency model and regenerating electrical energy.

### B. Robotic Exoskeleton

We used a research-grade robotic lower-limb exoskeleton (Exo-H3, Technaid) as an experimental platform to study energy regeneration and storage. The device weighs ∼12 kg and includes two leg modules, each with a thigh, shank, and foot segment, and a torso, which houses an onboard rechargeable lithium iron phosphate (LiFePO_4_) battery (22 V voltage at 12 Ah capacity) [18]. Six backdriveable actuators, each consisting of a harmonic drive and a wye-wound brushless DC motor, provide bilateral hip, knee, and ankle joint actuation in the sagittal-plane (i.e., six degrees of freedom in total). Each motor is rated at 19 V; has a terminal resistance and inductance phase-to-phase of 0.207 Ohms and 0.169 mH, respectively; a torque constant of 0.0375 Nm/A; a speed constant of 225 rpm/V; and a rotor inertia of 0.044 kg-cm^2^. The harmonic gearing has a transmission ratio of 160:1 and weighs ∼0.24 kg. The final peak torque output of motor-transmission system is 152 Nm with a reflected inertia of 8,346 kg-cm^2^ [18].

We used an external Controller Area Network (CAN) bus to communicate with the main onboard controller of the exoskeleton, which runs real-time control algorithms and interacts with the electronic drives of each motorized joint by acquiring sensor feedback and controlling the actuator. Each joint is equipped with an electronic drive board, which performs data acquisition of the onboard sensors, including those for joint angular position, interaction torque, and motor current (Fig. 3). The main controller and the individual joint controllers communicate at 1 Mbps using a real-time network based on CAN technology [18]. Each actuated joint can be controlled independently. The exoskeleton uses a hierarchical control architecture. The high-level controller includes separate controllers for different locomotor activities, which are manually selected using a mobile user interface. The low-level controller operates in either position control with prescribed joint kinematics, torque control, or stiffness control, which emulates a virtual spring system. According to the exoskeleton manufacturer, the actuator backdrive torque is ∼12 Nm, which is the minimum torque needed to overcome the mechanical impedance to backdrive the motor through its transmission.

**Figure 3.**
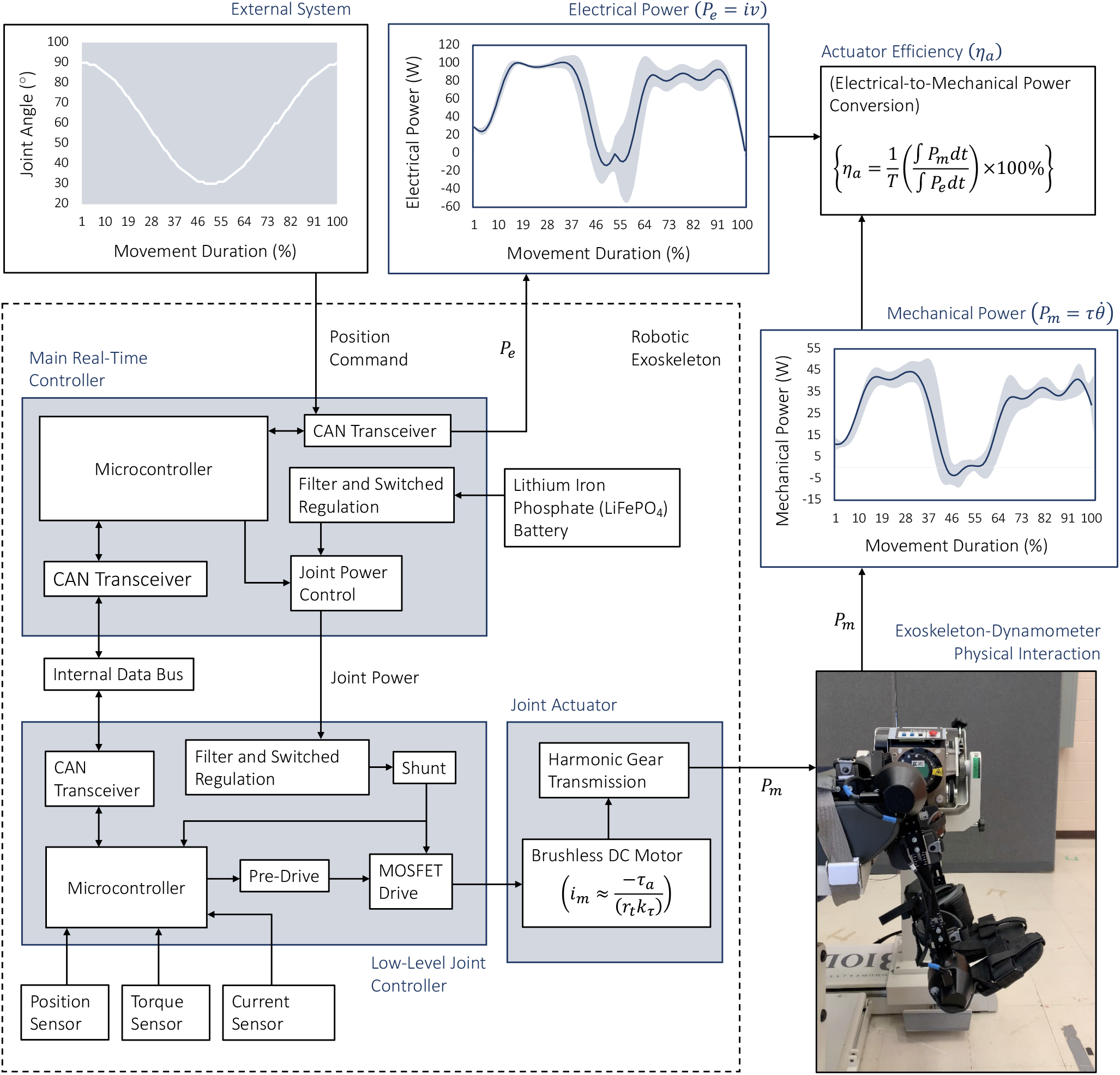
Exoskeleton benchtop testing with a joint dynamometer system to calculate actuator efficiency. The exoskeleton knee was kinematically driven through sinusoidal flexion and extension movements while the dynamometer measured the joint torque and angular velocity (i.e., the mechanical power output). The onboard exoskeleton sensors, and an electromechanical motor model, were used to estimate the motor voltage and current (i.e., the electrical power input). The actuator efficiency is the ratio of the average instantaneous power outputs to inputs over the steady-state time intervals. The nomenclature are described in the text. Details regarding the exoskeleton control architecture were taken from [18].

### C. Dynamometer Testing

We used a joint dynamometer system (Biodex) to measure the mechanical power output of the exoskeleton, similar to the setup in [16]. These dynamometers are commonly used in physical rehabilitation for isolated, single-joint testing to evaluate torque-angle (*τ* − *θ*) and torque-angular velocity (*τ* − *ω*) relationships [19] from which joint mechanical work and power can be calculated. The left exoskeleton knee was used for testing. Straps were used to secure the exoskeleton leg to the dynamometer attachment and the trunk segment to the seat. This helped prevent relative movement and misalignment between the rotational axes of the exoskeleton knee and the dynamometer shaft, which can cause errors in the torque measurements [19]. Using an upright seated posture, the exoskeleton knee was kinematically driven through 12-sinusoidal flexion and extension movements using position control (Fig. 3). The experiment was repeated four times with 10-minute breaks in between to prevent the exoskeleton motor from over-heating (n = 48 total trials). The isokinetic mode of the dynamometer was selected such that the exoskeleton could drive the shaft throughout the range of motion without interference since the controlled speed threshold far exceeded that of the exoskeleton joint.

During benchtop testing, the exoskeleton battery voltage and knee actuator torque were measured by onboard sensors, and the joint torque and angular velocity were measured by the dynamometer with a torque measurement accuracy of ± 1% of full scale (678 Nm). The dynamometer performed an automatic gravity correction on the measured joint torques by measuring the combined exoskeleton-dynamometer segment weight and applying the gravity correction based on the direction of shaft rotation. The weight was taken with the dynamometer such that the leg was positioned horizontally (i.e., maximum knee extension), wherein the gravity effect was highest, and a torque measurement was made. Depending on the exoskeleton leg orientation with respect to gravity during testing, the weight correction was either added or subtracted to the measured joint torques. Data were sampled at 100 Hz and filtered during postprocessing using a 10th-order low-pass Butterworth filter with an 8 Hz cut-off frequency and normalized to 0-100% of the movement duration. Since accelerations of the combined exoskeleton-dynamometer segment could produce unwanted inertial loads at the beginning and end of flexion and extension, only the middle portions of each movement, which were relatively constant speed, were used for data analyses (i.e., 15-35% and 65-85% of the overall sinusoidal movement).

### D. Actuator Efficiency

We calculated the exoskeleton actuator efficiency based on data from the dynamometer and onboard sensors, in addition to an electromechanical motor model. During standard motoring operation, the exoskeleton motor converts electrical power (*P*_*e*_) into mechanical power (*P*_*m*_). The mechanical power output is the product of the joint torque (*τ*_*j*_) and angular velocity 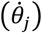 and the electrical power input is the product of the motor winding current (*i*_*m*_) and voltage (*𝒱*). When backdriven by an external load, the motor can operate like a generator, converting mechanical power into electrical power. The actuator efficiency (*η*_*a*_) during motoring operation is the ratio of electrical-to-mechanical power conversion 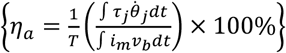 and vice-versa for energy regeneration when backdriven. The average instantaneous mechanical power output of the exoskeleton actuator was calculated as the product of the joint torque and angular velocity measured by the dynamometer 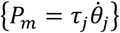 at each time step and averaged over the steady-state time intervals. Determining the average instantaneous electrical power input required additional consideration, as subsequently discussed.

Previous exoskeleton and wearable robotics studies [12], [20]–[22] have used the standard brushed DC electromechanical motor model governed by electrical and mechanical state equations. The motor winding voltage (*𝒱*_*m*_) can be mathematically modelled by applying Kirchhoff’s voltage law:

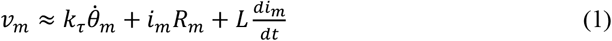

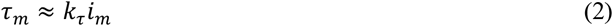

where *i*_*m*_ is the motor winding current, *k*_*τ*_ is the motor torque constant, *R*_*m*_ is the phase resistance of the motor windings, and *τ*_*m*_ and 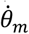 are the motor torque and angular velocity, respectively. The motor inductance 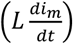 is relatively small and thus often omitted. The relationships between the torque-current and velocity-voltage data characterize the motor torque constant and back EMF constant, respectively; these constants are device-specific and depend on the motor topology and materials. Although the exoskeleton has onboard motor current sensors, the raw data are not available from the external CAN bus. However, the actuator torque (i.e., the combined motor-transmission system output) is provided. The motor phase current (*i*_*m*_) was thus back-calculated according to

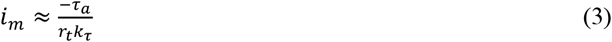

where *τ*_*a*_ is the actuator torque at each time step, *r*_*t*_ is the fixed transmission ratio of the harmonic gearing (160:1), and *k*_*τ*_ is the motor torque constant (0.0375 Nm/A) estimated by the motor manufacturer. The motor winding voltage (*𝒱*_*m*_) could not be directly solved for using equation (1) since the onboard exoskeleton sensors do not measure the motor angular velocity 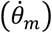 and the resolution of the joint angular position data was insufficient to back-calculate 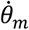 using the transmission ratio (*r*_*t*_). For simplicity, we assumed the measured battery voltage was equal to the voltage between the motor phases. The electrical power input used to drive the exoskeleton knee through flexion and extension was estimated by {*P*_*e*_ = *i*_*m*_*𝒱*_*b*_}, where *𝒱*_*b*_ is the measured battery voltage as reported by the external CAN bus. The reference axes of the motor variables and parameters were assumed to be equal since this information was not provided by the manufacturer. In other words, the three phase windings were assumed to be identical and a single set of parameters was used.

In theory, power losses (*P*_*loss*_) are mainly due to Joule heating, which is expressed by 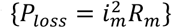. For example, in the MIT Cheetah robot, ∼76% of the energy dissipation was attributed to Joule heating [4]. Power losses from battery self-discharging and heating of the motor driver are relatively small and thus often ignored [21]. The actuator efficiency between trials (n = 48) was recorded to account for winding temperature changes over time. Although the exoskeleton is somewhat backdriveable, and thus theoretically capable of energy regeneration during negative mechanical work, the motor driver does not allow for electrical power flow back to the onboard battery. Therefore, we assumed that the power regeneration efficiency during backdrive operation was equal to the power generation efficiency during forward operation. There are ongoing discussions with the exoskeleton manufacturer as to how to safely and effectively regenerate energy without potential damage to the actuator, motor driver, and/or battery.

## III. Results

The human joint mechanical work, in addition to the calculated exoskeleton actuator efficiency from the benchtop testing (∼41%), were used to simulate power generation and regeneration. Appendix 1 shows the calculated hip, knee, and ankle joint mechanical work during human locomotion at variable speeds and slopes. Assuming a 75-kg user, the robotic exoskeleton could theoretically regenerate 6.3 J of total electrical energy per step during level-ground walking at 1 m/s since the corresponding total negative lower-limb joint mechanical work backdriving the actuators is 15.4 J/step. In comparison, walking downhill (e.g., -10° slope) at the same speed would require more negative joint mechanical work (27.8 J/step) for system deceleration and thus has greater potential for energy regeneration (11.4 J/step). Table 1 shows the estimated regenerated electrical energy for each combination of speed (0.8 m/s, 1 m/s, and 1.2 m/s) and slope (0° and ± 5° and 10°). Generally speaking, energy regeneration was inversely related to the incline slope but positively correlated to walking speed (i.e., faster walking down steeper slopes theorectically regenerates more electricity). Independent of speed and slope, the knee joint absorbed the most mechanical energy per step and thus has the greatest potential for energy regeneration.

**Table 1.** Estimated electrical energy regeneration (J) per step based on the exoskeleton actuator efficiency and negative joint mechanical work while walking at variable speeds (0.8 m/s, 1 m/s, and 1.2 m/s) and slopes (0° and ± 5° and 10°).

|  | -10° | -5° | 0° | 5° | 10° |
| --- | --- | --- | --- | --- | --- |
| 0.8 m/s | 10.0 | 7.0 | 5.4 | 3.4 | 2.8 |
| 1 m/s | 11.4 | 8.3 | 6.3 | 4.2 | 3.7 |
| 1.2 m/s | 12.8 | 9.4 | 7.4 | 5.3 | 4.7 |

Our performance calculations assume 1) regenerative braking from bilateral hip, knee, and ankle joints; 2) bidirectionally symmetric and constant (i.e., torque and velocity independent) exoskeleton actuator efficiencies; 3) power losses only from the motor-transmission system; 4) energy regeneration over the entire negative joint mechanical power range; and 5) identical actuator efficiencies at the hip, knee, and ankle joints. The exoskeleton and human joints were assumed to be kinematically constrained together, thereby ignoring any relative translations and rotations.

Integrating the positive joint mechanical power curves in Figs. 1 and 2 can provide insight into the electrical energy consumption during human-exoskeleton locomotion such that electrical energy from the onboard battery is used by the actuators to generate positive mechanical power. Table 2 shows the estimated electrical energy consumption for each combination of speed and slope. For example, level-ground walking at 1 m/s would require 21.4 J of total positive lower-limb joint mechanical work per step, therein equating to ∼52 J of total electrical energy consumption by the exoskeleton, assuming a 75-kg user and the actuator efficiency model. Using the parameter values of the lithium iron phosphate battery of the exoskeleton (22.4 V at 12 Ah for a total battery capacity of 967,680 J), the device could theoretically walk 18,624 steps per battery charge on level-ground at 1 m/s (Table 3). In comparison, with energy regeneration, the battery-powered operating time could be extended by an additional 14% (i.e., 2,575 additional steps for 21,199 total steps (*T*_*steps*_)) such that

**Table 2.**
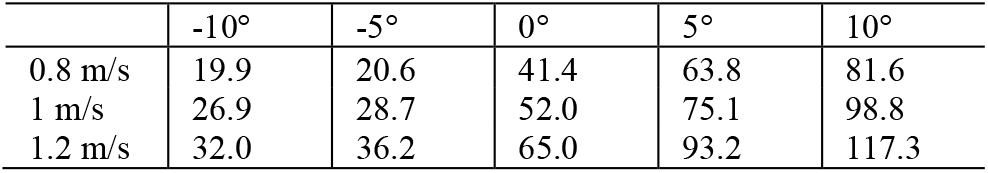
Estimated electrical energy consumption (J) per step based on the exoskeleton actuator efficiency and positive joint mechanical work while walking at variable speeds (0.8 m/s, 1 m/s, and 1.2 m/s) and slopes (0° and ± 5° and 10°).

**Table 3.** Estimated number of steps per battery charge while walking at variable speeds (0.8 m/s, 1 m/s, and 1.2 m/s) and slopes (0° and ± 5° and 10°) *without energy regeneration*. Calculations were based on the exoskeleton battery parameters and results in Table 2.

|  | -10° | -5° | 0° | 5° | 10° |
| --- | --- | --- | --- | --- | --- |
| 0.8 m/s | 48,533 | 47,048 | 23,400 | 15,177 | 11,858 |
| 1 m/s | 35,987 | 33,726 | 18,624 | 12,892 | 9,795 |
| 1.2 m/s | 30,260 | 26,734 | 14,898 | 10,382 | 8,248 |

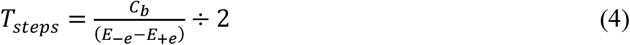

where *C*_*b*_ is the total capacity of the lithium iron phosphate battery, *E*_−*e*_ is the total electrical energy consumption per step by the exoskeleton, and *E*_+*e*_ is the total electrical energy that could theoretically be regenerated per step. Here the negative and positive signs represent the electrical power flowing in and out of the battery during power generation and regeneration, respectively. These performance calculations assume an adequate motor driver to control the bidirectional flow of electrical power between the motor and onboard battery. The electrical-to-chemical energy conversion efficiency of the battery was ignored.

Table 4 shows the estimated number of steps per battery charge with energy regeneration for each speed and slope combination. The greatest performance benefits were observed while walking at the slowest speed (0.8 m/s) and steepest decline (−10°), where the battery-powered operating time could theorectically extend by 99% compared to not regenerating energy. Although negative joint mechanical work increases with walking speed, and thus absorbs more energy and increases the potential for energy regeneration, the positive joint mechanical work used for forward propulsion, and thus the electrical power consumption by the exoskeleton, disproportionally increases. Therefore, the locomotor efficiency is theorectically greater at slower walking speeds due to the ratio of negative-to-positive joint mechanical work, therein yielding longer battery-powered operating times.

**Table 4.**
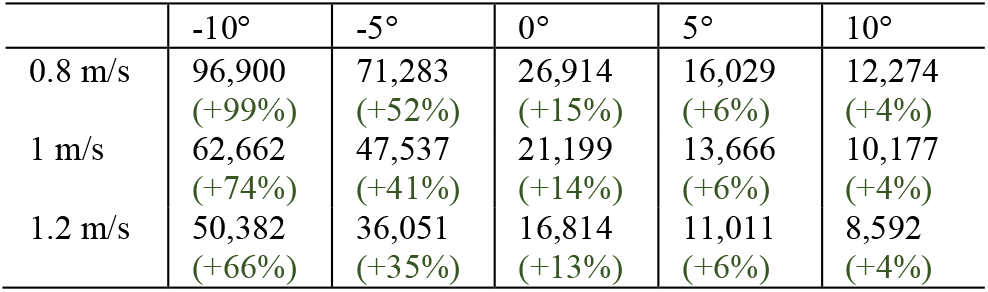
Estimated number of steps per battery charge while walking at variable speeds (0.8 m/s, 1 m/s, and 1.2 m/s) and slopes (0° and ± 5° and 10°) *with energy regeneration*. The percent (%) increase in step count is also provided. Calculations were based on equation (4).

|  | -10° | -5° | 0° | 5° | 10° |
| --- | --- | --- | --- | --- | --- |
| 0.8 m/s | 96,900<br>(+99%) | 71,283<br>(+52%) | 26,914<br>(+15%) | 16,029<br>(+6%) | 12,274<br>(+4%) |
| 1 m/s | 62,662<br>(+74%) | 47,537<br>(+41%) | 21,199<br>(+14%) | 13,666<br>(+6%) | 10,177<br>(+4%) |
| 1.2 m/s | 50,382<br>(+66%) | 36,051<br>(+35%) | 16,814<br>(+13%) | 11,011<br>(+6%) | 8,592<br>(+4%) |

## IV. Discussion

In this study, we present the development of a feedforward human-exoskeleton energy regeneration system model to simulate energy regeneration and storage during daily locomotor activities. Data from inverse dynamics analyses [17] of walking at variable speeds and slopes were used to calculate negative joint mechanical power and work (i.e., the mechanical energy theoretically available for electrical energy regeneration). These human joint mechanical energetics were then used to simulate backdriving an exoskeleton and regenerating energy. An empirical characterization of the exoskeleton device was carried out using a joint dynamometer system and an electro-mechanical motor model to calculate the actuator efficiency and to simulate energy regeneration. Our performance calculations showed that regenerating energy at slower walking speeds and decline slopes could significantly extend the battery-powered operating times of robotic lower-limb exoskeletons (i.e., up to 99% increase in total number of steps), therein improving locomotor efficiency. In addition to exoskeletons, these principles of energy regeneration and storage could extend to robotic leg prostheses, powered wheelchairs, and other mobility assistive devices.

Our calculations also showed that, independent of speed and slope, the knee absorbs the most mechanical energy during human locomotion and thus has the greatest potential for energy regeneration. This differs from our previous simulation-based research on human-exoskeleton stand-to-sit movements, which showed that the hip undergoes the most negative joint mechanical work [23]. Most exoskeletons with regenerative actuators have focused on knee designs [6], [11], [12], [21]. In addition to having more favorable mechanical energetics, the knee is typically preferred for energy regeneration since 1) the hip is more structurally complex, and 2) many wearable robotic devices, especially prosthetic legs, do not include a hip joint. For optimal efficiency and battery performance, robotic exoskeletons should be designed to recover the otherwise dissipated energy at each motorized joint during periods of negative mechanical work.

One of the biggest limitations to energy regeneration is the relatively low efficiency of most motor-transmission systems. The calculated exoskeleton actuator efficiency from our benchtop testing was ∼41% during forward operation (i.e., electrical-to-mechanical power conversion); we assumed bidirectional symmetry of the actuator efficiency to simulate energy regeneration when backdriven. In comparison, the MIT Cheetah robot [4] and the biomechanical energy harvesting knee exoskeleton by Donelan and colleagues [11], [12] both achieved ∼63% actuator efficiency during mechanical-to-electrical power conversion. To our knowledge, these energy regeneration efficiencies are some of the highest reported in the literature for legged systems.

Two of the leading sources of energy losses are Joule heating in the motor windings and friction in the transmission [2], [5]. High transmission ratios can reduce the motor torque needed for legged locomotion, and thus decrease the motor current and Joule heating losses. However, high gearing tends to increase the actuator weight, friction, and reflected inertia, which increase impedance and decreases backdrivability and the potential for energy regeneration [2]. Alternatively, high torque-density motors can decrease the needed transmission ratios by generating high output torques, thus circumventing the inefficiencies of high gearing, although at the expense of more motor current and higher Joule heating losses [5]. For example, ∼76% of the energy losses in the MIT Cheetah were attributed to Joule heating, which was designed with quasi-direct drives (6:1 ratio with planetary gearing) [4]. Note that the selection of gearing, in addition to increasing the gear ratio, can increase the energy losses due to friction.

An open challenge for the research community is to optimize the tradeoff between actuator output torque and backdrive torque in terms of efficiency (including energy regeneration) and performance. However, given the complex interactions between different actuator design parameters, determining the optimal system design via experimental trial-and-error would be difficult. Modelling and simulation of an integrated human-exoskeleton system [16] could allow for efficient testing and co-optimization of different actuator design parameters while also taking into consideration changes in human biomechanics and metabolic energetics. This differs from our study, which used independent models of the human and exoskeleton systems without including closed-loop interactions. This a computational framework with a higher-fidelity actuator model could help address some of the simplifying assumptions made in our performance calculations such as bidirectionally symmetric and constant (i.e., torque and velocity independent) actuator efficiencies.

Lastly, our performance calculations were based on data from healthy young adults [17], thus requiring several assumptions and extrapolations to aging and rehabilitation populations. Healthy young adults typically walk ∼1.4 m/s and take 6,000-13,000 steps/day [24]. However, persons with mobility impairments tend to walk slower and take fewer steps per day. For example, the average self-selected walking speed in adults over 75 years age is ∼24% slower compared to those 25 years age [24]. The joint energetics during locomotion also differ between these populations [25]. Slower walking speeds have implications on energy regeneration since faster walking tends to generate more electricity and more efficiently [12]. Slower walking would backdrive the actuator with lower rotational speeds and thus generate less electricity. Motors are also typically less efficient when generating torques at low speeds due to Joule heating [5]. Given these biomechanical and activity level differences between healthy young adults and persons with mobility impairments, and the implications of such differences on the actuator performance, future research should study older adults and/or those with physical disabilities to improve our energy regeneration performance calculations.

## Acknowledgment

We acknowledge Elkyn Belalcazar for his assistance with the exoskeleton benchtop testing. We thank Robert Gregg and Emma Reznick for their support with the biomechanics dataset. We also thank Ung Hee Lee and Edgar Bolívar for their discussions on wearable robotic actuators.

---

**Appendix 1.** Hip, knee, and ankle joint mechanical work (J/kg) per stride in healthy young adults walking at variable speeds (0.8 m/s, 1 m/s, and 1.2 m/s) and slopes (0° and ± 5° and 10°). The results are averages across multiple subjects (n=10) and strides and normalized to total body mass. Data were calculated from [17]. “Total Work” is the combined mechanical energies from the hip, knee, and ankle joints and “Net Joint Work” is the net mechanical work on each joint. The total negative lower-limb joint mechanical work represents the amount of mechanical energy theoretically available for electrical energy regeneration. The results are for individual legs and are reported for (−10°, -5°, 0°, +5°, +10°) slopes.

|  | Speed (m/s) | Positive Work (J/kg) | Negative Work (J/kg) | Net Joint Work (J/kg) |
| --- | --- | --- | --- | --- |
| Hip | 0.8 | (0.09, 0.09, 0.20, 0.32, 0.44) | (-0.03, -0.04, -0.02, -0.03, -0.03) | (0.06, 0.05, 0.18, 0.29, 0.41) |
|  | 1 | (0.12, 0.13, 0.24, 0.38, 0.56) | (-0.06, -0.06, -0.03, -0.04, -0.04) | (0.06, 0.07, 0.21, 0.34, 0.51) |
|  | 1.2 | (0.15, 0.16, 0.30, 0.48, 0.63) | (-0.09, -0.08, -0.05, -0.06, -0.06) | (0.06, 0.07, 0.25, 0.42, 0.57) |
| Knee | 0.8 | (0.03, 0.02, 0.06, 0.10, 0.13) | (-0.37, -0.23, -0.16, -0.14, -0.12) | (-0.34, -0.20, -0.09, -0.04, 0.01) |
|  | 1 | (0.05, 0.04, 0.09, 0.12, 0.15) | (-0.45, -0.29, -0.22, -0.18, -0.17) | (-0.40, -0.25, -0.13, -0.07, -0.02) |
|  | 1.2 | (0.06, 0.07, 0.12, 0.14, 0.16) | (-0.51, -0.35, -0.28, -0.25, -0.22) | (-0.44, -0.28, -0.16, -0.10, -0.06) |
| Ankle | 0.8 | (0.10, 0.11, 0.19, 0.28, 0.32) | (-0.25, -0.19, -0.18, -0.05, -0.03) | (-0.15, -0.08, 0.01, 0.23, 0.29) |
|  | 1 | (0.13, 0.14, 0.24, 0.32, 0.37) | (-0.24, -0.19, -0.16, -0.05, -0.03) | (-0.11, -0.05, 0.07, 0.27, 0.34) |
|  | 1.2 | (0.14, 0.17, 0.29, 0.40, 0.49) | (-0.23, -0.17, -0.15, -0.04, -0.03) | (-0.09, 0.00, 0.14, 0.36, 0.47) |
| Total Work | 0.8 | (0.22, 0.22, 0.45, 0.70, 0.89) | (-0.65, -0.45, -0.35, -0.22, -0.18) | (-0.43, -0.23, 0.10, 0.48, 0.71) |
|  | 1 | (0.29, 0.31, 0.57, 0.82, 1.08) | (-0.74, -0.54, -0.41, -0.28, -0.24) | (-0.45, -0.23, 0.16, 0.54, 0.84) |
|  | 1.2 | (0.35, 0.40, 0.71, 1.02, 1.28) | (-0.83, -0.61, -0.48, -0.35, -0.31) | (-0.48, -0.21, 0.23, 0.67, 0.98) |

## References

[1] A. J. Young and D. P. Ferris, “State of the art and future directions for lower limb robotic exoskeletons,” IEEE Trans. Neural Syst. Rehabil. Eng., vol. 25, no. 2, pp. 171–182, Feb. 2017.

[2] P. L. García, S. Crispel, E. Saerens, T. Verstraten, and D. Lefeber, “Compact gearboxes for modern robotics: A review,” Front. Robot. AI, vol. 7, p. 103, Aug. 2020.

[3] N. Kashiri et al., “An overview on principles for energy efficient robot locomotion,” Front. Robot. AI, vol. 5, p. 129, Dec. 2018.

[4] S. Seok et al., “Design principles for energy-efficient legged locomotion and implementation on the MIT cheetah robot,” IEEE/ASME Trans. Mechatron., vol. 20, no. 3, pp. 1117–1129, Jun. 2015.

[5] J. Hollerbach, I. Hunter, and J. Ballantyne, “A comparative analysis of actuator technologies for robotics,” in The Robotics Review 2, 1992, pp. 299–342.

[6] B. Laschowski, J. McPhee, and J. Andrysek, “Lower-limb prostheses and exoskeletons with energy regeneration: Mechatronic design and optimization review,” ASME J. Mech. Robot., vol. 11, no. 4, p. 040801, Aug. 2019.

[7] H. Zhu, C. Nesler, N. Divekar, M. T. Ahmad, and R. D. Gregg, “Design and validation of a partial-assist knee orthosis with compact, backdrivable actuation,” in 2019 IEEE 16th International Conference on Rehabilitation Robotics (ICORR), Toronto, ON, Canada, Jun. 2019, pp. 917–924.

[8] G. Lv, H. Zhu, and R. D. Gregg, “On the design and control of highly backdrivable lower-limb exoskeletons: A discussion of past and ongoing work,” IEEE Control Syst., vol. 38, no. 6, pp. 88–113, Dec. 2018.

[9] H. Zhu, C. Nesler, N. Divekar, V. Peddinti, and R. Gregg, “Design principles for compact, backdrivable actuation in partial-assist powered knee orthoses,” IEEE/ASME Trans. Mechatron., pp. 1–1, 2021.

[10] C. Nesler, G. Thomas, N. Divekar, E. J. Rouse, and R. D. Gregg, “Enhancing voluntary motion with modular, backdrivable, powered hip and knee orthoses,” IEEE Robot. Autom. Lett., Jan. 2022.

[11] J. M. Donelan, Q. Li, V. Naing, J. A. Hoffer, D. J. Weber, and A. D. Kuo, “Biomechanical energy harvesting: Generating electricity during walking with minimal user effort,” Science, vol. 319, no. 5864, pp. 807–810, Feb. 2008.

[12] Q. Li, V. Naing, and J. M. Donelan, “Development of a biomechanical energy harvester,” J. NeuroEngineering Rehabil., vol. 6, no. 1, p. 22, Dec. 2009.

[13] M. S. Orendurff, J. Schoen, G. Bernatz, A. Segal, and G. Klute, “How humans walk: Bout duration, steps per bout, and rest duration,” J. Rehabil. Res. Dev., vol. 45, no. 7, pp. 1077–1090, Dec. 2008.

[14] B. Laschowski, W. McNally, A. Wong, and J. McPhee, “ExoNet database: Wearable camera images of human locomotion environments,” Front. Robot. AI, vol. 7, p. 562061, Dec. 2020.

[15] B. Laschowski, “Energy regeneration and environment sensing for robotic leg prostheses and exoskeletons,” PhD Thesis, University of Waterloo, 2021.

[16] K. A. Inkol and J. McPhee, “Assessing control of fixed-support balance recovery in wearable lower-limb exoskeletons using multibody dynamic modelling,” in 2020 8th IEEE RAS/EMBS International Conference for Biomedical Robotics and Biomechatronics (BioRob), New York City, NY, USA, Nov. 2020, pp. 54–60.

[17] E. Reznick, K. R. Embry, R. Neuman, E. Bolívar-Nieto, N. P. Fey, and R. D. Gregg, “Lower-limb kinematics and kinetics during continuously varying human locomotion,” Sci. Data, vol. 8, no. 1, p. 282, Dec. 2021.

[18] M. Bortole et al., “The H2 robotic exoskeleton for gait rehabilitation after stroke: Early findings from a clinical study,” J. NeuroEngineering Rehabil., vol. 12, no. 1, p. 54, Dec. 2015.

[19] W. Herzog, “The relation between the resultant moments at a joint and the moments measured by an isokinetic dynamometer,” J. Biomech., vol. 21, no. 1, pp. 5–12, Jan. 1988.

[20] U. H. Lee, C.-W. Pan, and E. J. Rouse, “Empirical characterization of a high-performance exterior-rotor type brushless DC motor and drive,” in 2019 IEEE/RSJ International Conference on Intelligent Robots and Systems (IROS), Macau, China, Nov. 2019, pp. 8018–8025.

[21] E. Bolívar, S. Rezazadeh, and R. Gregg, “A general framework for minimizing energy consumption of series elastic actuators with regeneration,” in ASME 2017 Dynamic Systems and Control Conference, Tysons, Virginia, USA, Oct. 2017, p. V001T36A005.

[22] H. Richter, “A framework for control of robots with energy regeneration,” ASME J. Dyn. Syst. Meas. Control, vol. 137, no. 9, p. 091004, Sep. 2015.

[23] B. Laschowski, R. S. Razavian, and J. McPhee, “Simulation of stand-to-sit biomechanics for robotic exoskeletons and prostheses with energy regeneration,” IEEE Trans. Med. Robot. Bionics, vol. 3, no. 2, pp. 455–462, May 2021.

[24] M. Grimmer, R. Riener, C. J. Walsh, and A. Seyfarth, “Mobility related physical and functional losses due to aging and disease -A motivation for lower limb exoskeletons,” J. NeuroEngineering Rehabil., vol. 16, no. 1, p. 2, Jan. 2019.

[25] D. A. Winter, The Biomechanics and Motor Control of Human Gait: Normal, Elderly and Pathological, 2nd Edition. Waterloo, Canada: Waterloo Biomechanics, 1991.

